# Role of protonation states in stability of molecular dynamics simulations of high-resolution membrane protein structures

**DOI:** 10.1101/2023.08.24.554589

**Authors:** Jonathan Lasham, Amina Djurabekova, Volker Zickermann, Janet Vonck, Vivek Sharma

## Abstract

Classical molecular dynamics (MD) simulations provide unmatched spatial and time resolution of protein structure and function. However, accuracy of MD simulations often depends on the quality of force field parameters and the time scale of sampling. Another limitation of conventional MD simulations is that the protonation states of titratable amino acid residues remain fixed during simulations, even though protonation state changes coupled to conformational dynamics are central to protein function. Due to the uncertainty in selecting protonation states, classical MD simulations are sometimes performed with all amino acids modeled in their standard charged states at pH 7. Here we performed and analyzed classical MD simulations on high-resolution cryo-EM structures of two membrane proteins that transfer protons by catalyzing protonation/deprotonation reactions. In simulations performed with amino acids modeled in their standard protonation state the structure diverges far from its starting conformation. In comparison, MD simulations performed with pre-determined protonation states of amino acid residues reproduce the structural conformation, protein hydration, and protein-water and protein-protein interactions of the structure much better. The results suggest it is crucial to perform basic protonation state calculations, especially on structures where protonation changes play an important functional role, prior to launching any MD simulations. Furthermore, the combined approach of protonation state prediction and MD simulations can provide valuable information on the charge states of amino acids in the cryo-EM sample. Even though accurate prediction of protonation states currently remains a challenge, we introduce an approach of combining pKa prediction with cryo-EM density map analysis that helps in improving not only the protonation state predictions, but also the atomic modeling of density data.

## Introduction

Within the past decade, the resolution of single-particle cryo-EM structures has improved dramatically, largely due to the improvements in direct electron detectors and processing software ^1^. The resolution of single-particle cryo-EM structures is now comparable to x-ray crystallography and NMR structures, and the so-called “resolution revolution” has made it possible to determine structures of many previously inaccessible complexes, particularly for membrane proteins ^2^. Many of these structures have a resolution below 2.5 Å, allowing accurate modeling of protein conformations, including ordered water molecules, which has significant implications in drug design ^3^ and is also central to our understanding of the molecular mechanism of enzymes that catalyze proton transfer reactions ^4^.

Proton transfer can take place across a chain of water molecules via a Grotthuss-type mechanism ^5^. However, proton transfer routes through proteins do not consist only of water molecules but are often made up of hydrogen-bonded networks of polar and charged amino acid residues. Besides titratable residues such as glutamic acid and lysine, which can donate or accept protons, other residues that are part of these networks include asparagine and glutamine, and serine, threonine, and tyrosine with hydroxyl groups in their sidechains. These polar residues can not only assist in proton transfer reactions by stabilizing charged intermediates but may also undergo protonation/deprotonation reactions in proteinaceous environments ^4, 6–8^. Moreover, titratable acidic and basic residues are well-known to act as proton transfer elements and proton loading sites in several enzymes ^4, 9, 10^. However, the protonation states in protein structures are usually not explicitly modelled. Such information can be obtained with neutron diffraction techniques, but only for relatively small proteins ^11^.

The two main structure determination methods for protein complexes, x-ray crystallography and cryo-EM, produce superficially similar results. However, they differ fundamentally in the way atomic structures are imaged. X-rays are deflected by electrons and thus produce electron density maps, but electrons are scattered by Coulomb potential, and they are thus sensitive to charges. While positively charged ions just add extra density to an already positive signal, negative charges, in particular O^-^, can give rise to negative electron scattering amplitudes ^12–14^. As a result, negatively charged sidechains of glutamate and aspartate residues are not visible in cryo-EM maps. Although the weak density of these sidechains has sometimes been interpreted as a result of the radiation sensitivity of the carboxyl group ^15, 16^, the former mechanism appears to have a larger contribution ^14, 17–19^. The absence of sidechain density hampers accurate model building. On the other hand, it makes it possible to determine the charge state of acidic residues.

To complement cryo-EM structures, molecular dynamics (MD) simulations are often performed ^20–30^. These help in studying the dynamics of the protein and the solvent, as well as the binding and unbinding of lipids, ligands, and ions. The approach of combining structural data with MD simulations is a powerful technique to understand protein structure and function. The abundance of new structures gives plenty of opportunity to perform these simulations routinely at various levels of computational approximations. However, one limitation of conventional molecular dynamics is that covalent bonds are fixed throughout the simulation, therefore the charge states of titratable residues (often selected as the standard charge state that is Asp/Glu deprotonated, Lys/Arg protonated and His neutral) remain unchanged. This results in a biased scenario, and biologically relevant conformational states with alternative protonation states are not populated. Moreover, situations where changes in protonation states can occur as a function of conformational dynamics (e.g. membrane proteins catalyzing proton transfer) are also not captured. However, there are various methods that can allow charge states of amino acids to change during MD simulations, such as hybrid QM/MM MD, and constant pH MD, but these are generally computationally costly ^31^, although there have been recent improvements in the enhanced scalability of constant pH MD ^32^.

For many proteins, performing MD simulations by systematic altering of charge states of individual titratable residues is unrealistic, due to the large number of such residues. Of course, one can alter protonation states of selected conserved residues to study specific questions, and this has indeed been performed to obtain valuable functional insights ^33–37^. However, due to the long-range nature of electrostatic interactions the protonation state of one titratable residue affects another’s by ca. 8 kcal/mol (with a separation of 10 Å at ε=4) and such aspects are often ignored when performing simulations.

Alternatively, one can calculate the pKa of all titratable residues in a protein for a given conformational state (obtained from a cryo-EM experiment, for example) and perform MD simulations in that fixed protonation state. There are several methods to perform pKa calculations, many of which are based on continuum electrostatics approaches ^38^. However, these require a significant amount of preprocessing and can display large variation in pKa values due to subtle conformational changes. Alternatively, empirical methods, such as Propka can be employed relatively easily, to give fast and sufficiently accurate predictions of the pKa of all ionizable groups present in a protein, even if hundreds in number ^39^. Methods like Propka ^40^, often give reasonable estimates for the pKa of buried titratable residues, in particular if the sites are far from any redox active cofactors. Due to their rapid pKa estimations, they can be used on thousands of simulation snapshots to obtain profiles of pKa change as a function of conformational dynamics ^35, 41^. Similarly, Monte Carlo-based pKa prediction methods have also been used to predict charged states of systems, either using standalone PDB files or simulation snapshots ^42^.

In this study, we present MD simulations on two high-resolution cryo-EM structures of membrane proteins. The two structures are the respiratory complex I from *Yarrowia lipolytica* (PDB 7O71, EMD-12742) ^43^ and the multiple resistance and pH adaptation (Mrp) cation/proton antiporter from *Bacillus pseudofirmus* (PDB 7QRU, EMD-14124) ^44^. Both proteins facilitate proton transfer reactions that involve amino acid residues and water molecules, many of which have been resolved in the two structures. The role of protein hydration, water dynamics as well as change in protonation state is central to the structure and function of these proteins. By performing long time scale MD simulations in multiple charge states of these proteins and extensively analyzing protein and solvent dynamics, we show that both proteins deviate from the original cryo-EM conformation when simulated in the standard protonation state of titratable amino acid residues, while MD simulations in pre-defined protonation states stabilize the protein conformation much better. We propose that the approach of protonation state prediction combined with MD simulations can give insights into the charge state of residues in a cryo-EM structure. Furthermore, we find that the prediction of protonation states of acidic residues agrees well with the charge state assignment based on cryo-EM density maps, and that outliers can be identified with the approach discussed here, leading to an improved modeling of cryo-EM density data.

## Results

Two high-resolution membrane protein structures were chosen for this investigation. First, the 2.1 Å resolution structure of complex I from *Y. lipolytica*, which is over 1 MDa in size, and has more than 1600 structural water molecules resolved ^43^. Around 100 of these are in the potential proton transfer pathways present in its membrane-bound subunits (Fig. 1). Second, a structure from the evolutionarily related Na^+^/H^+^ Mrp antiporter ^44^ at a similar resolution of 2.2 Å. This protein is much smaller than complex I (ca. 213 kDa) and consists of membrane-bound subunits only, without redox active cofactors. There are 360 water molecules resolved in the structure of the Mrp antiporter, around 70 of them in the potential proton and sodium transfer pathways (Fig. 1). The structure of complex I contains more than 1300 titratable residues, while the Mrp structure has around 200. We performed MD simulations on both complexes with all titratable sites either modeled in their standard states (S state) or based on pKa calculations at pH 7 (P7 state). We find that in the P7 state, complex I has 68 neutralized titratable amino acids, out of which 25 are in the membrane core of the enzyme (Fig. 1), while in the Mrp antiporter 13 residues are neutralized in the P7 state. Accordingly, the charge reduces from +90/-94 to +83/-76 in the complex I membrane arm, and from +116/- 118 to +110/-112 in the Mrp upon neutralization of titratable sites as part of pKa calculations (see Tables S1 and S2 for lists of residues neutralized in complex I and Mrp antiporter, respectively).

**Fig. 1.**
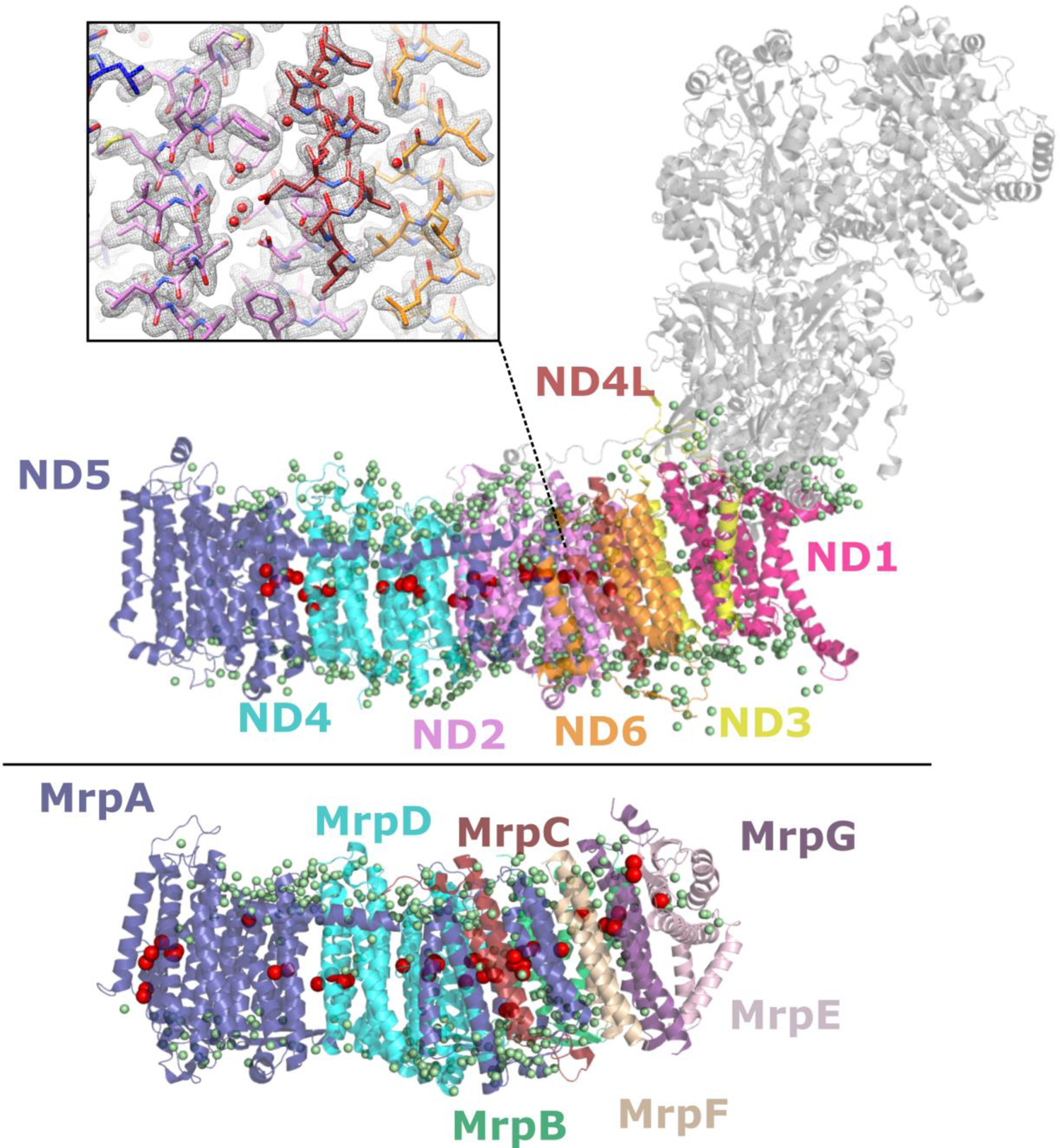
Structure of the membrane domain of respiratory complex I (upper) and Mrp antiporter (lower). The protein is shown in ribbon representation and colored by subunit. Only core membrane-bound subunits are shown in color for complex I, with the hydrophilic core subunits shown as gray. Structurally resolved water molecules are shown as small green spheres, with those in the functionally relevant hydrophilic axes of complex I and Mrp antiporter shown as larger red spheres. Proton translocation is suggested to take place across this axis, which is a vital part of the mechanism for both proteins. The inset in the upper panel shows a region of the complex I membrane arm within the high-resolution cryo-EM density (grey mesh), including density for water molecules.

### Global mobility of proteins in P7 and S states

First, the overall global mobility of both systems was analyzed in the S and P7 states. Fig. 2 shows the time series as well as the distribution of the RMSD (root mean square deviation) of the backbone atoms for the core membrane-bound subunits of complex I and Mrp antiporter in the two simulation states. The P7 state systems with lower charge have overall lower RMSD for both protein complexes, which means that the backbone atoms stay closer to the starting conformation throughout the simulation. This notion is also true if a similar RMSD analysis is performed on the Cα backbone atoms and all protein atoms excluding hydrogens, even with the inclusion of the hydrophilic domain of complex I (Fig. S1).

**Fig. 2.**
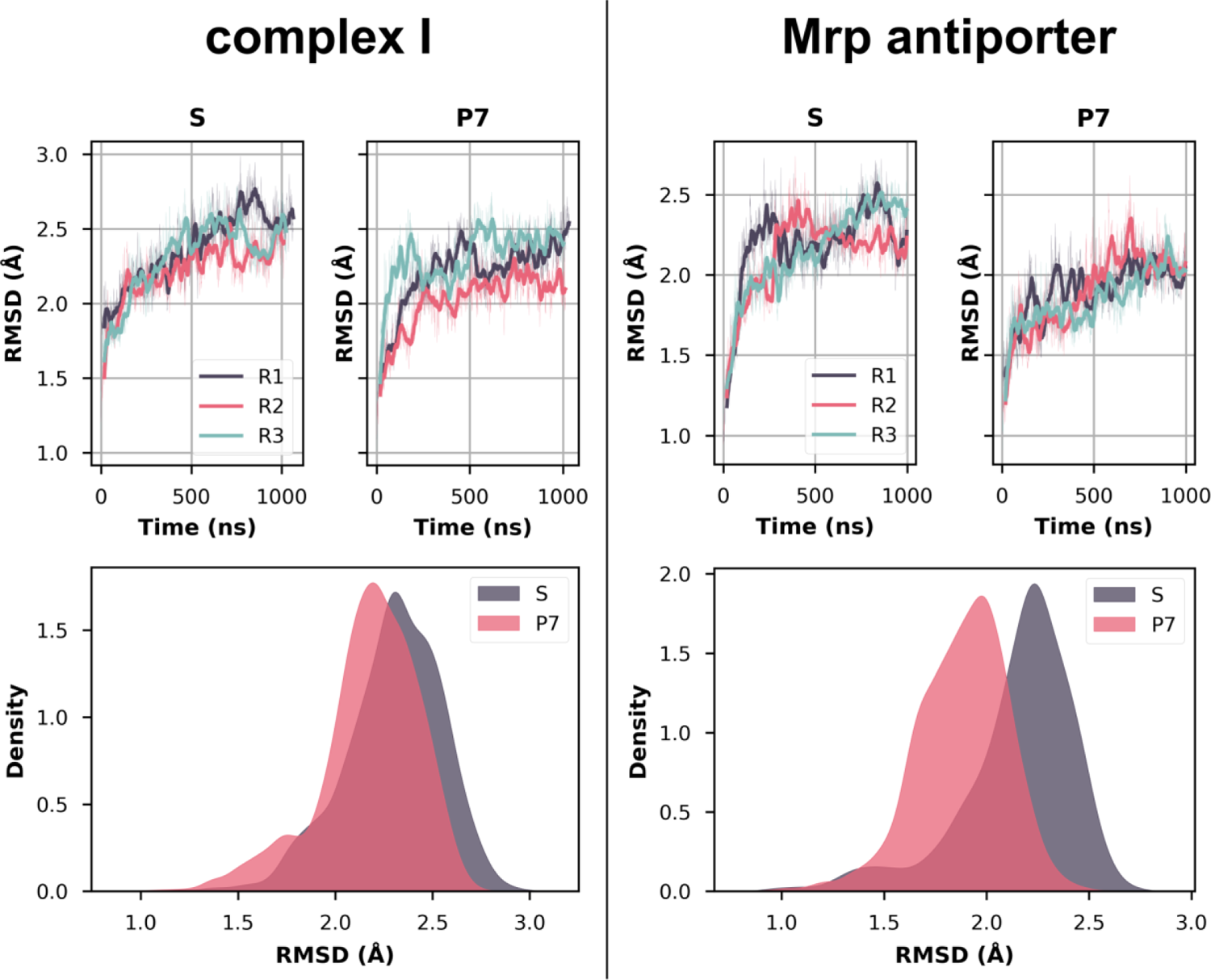
RMSD (root mean square deviation) of the backbone atoms over time for both complex I (core membrane subunits, left panel) and Mrp antiporter (right panel) in S and P7 states. The upper plots show the RMSD as a time series, with each colored trace representing a different simulation replica. The thick lines show the moving average of 20 ns, while the thinner lines show the RMSD for every 1 ns. The bottom panels show the distribution of RMSD values in the S and P7 states using a kernel density estimate (KDE) function with combined data of all three replicas.

Similarly, the RMSF (root mean square fluctuation), which measures the average amount an atom moves during the entire simulation, is also consistently found to be lower in the P7 states than in the S states. The boxplots in Fig. 3 show the RMSF of Cα atoms in each individual subunit of the Mrp antiporter. For all antiporter chains, lower RMSF values are observed in the P7 state, indicating that there is an overall stability in the system as charges of titratable sites are neutralized (based on pKa estimates). A similar trend was also observed in complex I (Fig. S2). The spatial distribution of the RMSF data is shown in the lower panel of Fig. 3. MrpA subunit (Figs. 1 and 3) has the largest number of titratable residues that undergo protonation state changes to become neutralized and this has a clear impact in reducing the RMSF of the subunit. Interestingly, the stability of the protein also increases in areas somewhat far from the residues that change protonation states (magenta spheres), highlighting that the charge change of just a few titratable residues can have conformational changes imparted both at the local and the global level.

**Fig. 3.**
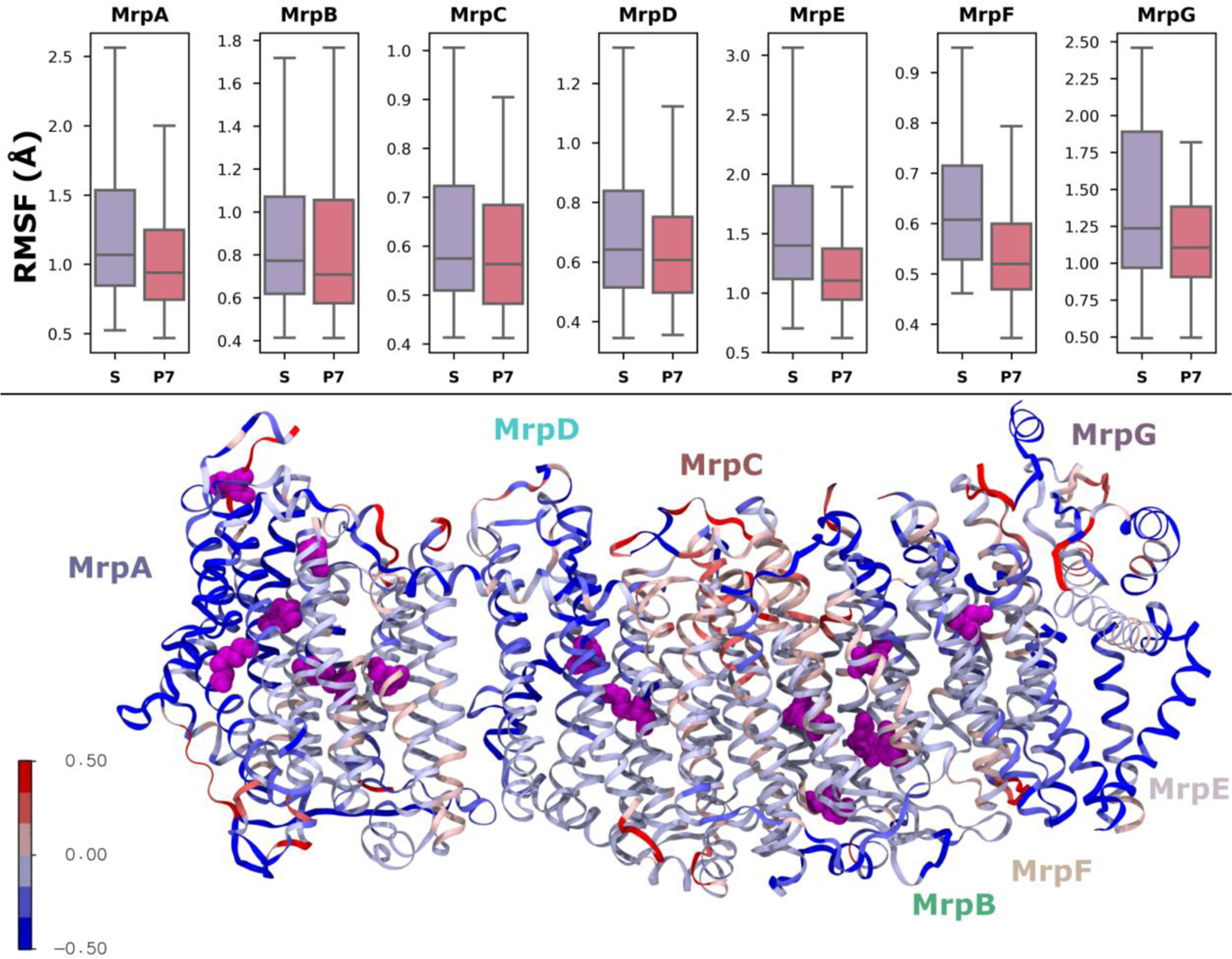
(Top) RMSF (root mean square fluctuation) of Cα atoms in Mrp antiporter subunits, calculated for all simulation data in S and P7 states. The box plots represent the distribution of RMSF values for each subunit, in both states. The shaded box represents the interquartile range, with the middle line showing the median. The upper and lower lines are the maximum and minimum values respectively. (Bottom) The panel shows the protein structure colored according to the extent of conformational change in the P7 state compared to the S state. Red represents Cα RMSF increased in the P7 state and blue represents a decrease. The magenta spheres show the positions of residues that undergo a protonation state change in P7 simulations.

Overall, the key observation is that MD simulations performed in predefined protonation states retain a structure closer to the cryo-EM conformation for both protein complexes studied here. This in turn means that the simulated charge state is closer to the charge state of the protein during cryo-EM sample preparation. To understand why P7 state simulations show overall stability relative to the S state, we performed additional analyses.

### Water-protein interactions and protein hydration in P7 and S states

In the high-resolution structures of complex I and Mrp antiporter, the positions of several water molecules are resolved. We therefore next analyzed how water-protein interactions and protein hydration are affected in the S and P7 simulation states. The contacts between all water molecules and protein were clearly reduced in the P7 states compared with the S states (Fig. 3, upper panels). The lower charge in P7 states (+83/-76 compared to +90/-94 in S state of complex I and +110/-112 compared to +116/-118 in S state of Mrp antiporter) prevents extensive hydration of the protein resulting in lower number of contacts between water oxygens and protein. This is also reflected in the clear reduction of water contacts that take place with the residues that change protonation state (Fig 3, middle panel).

**Fig. 3.**
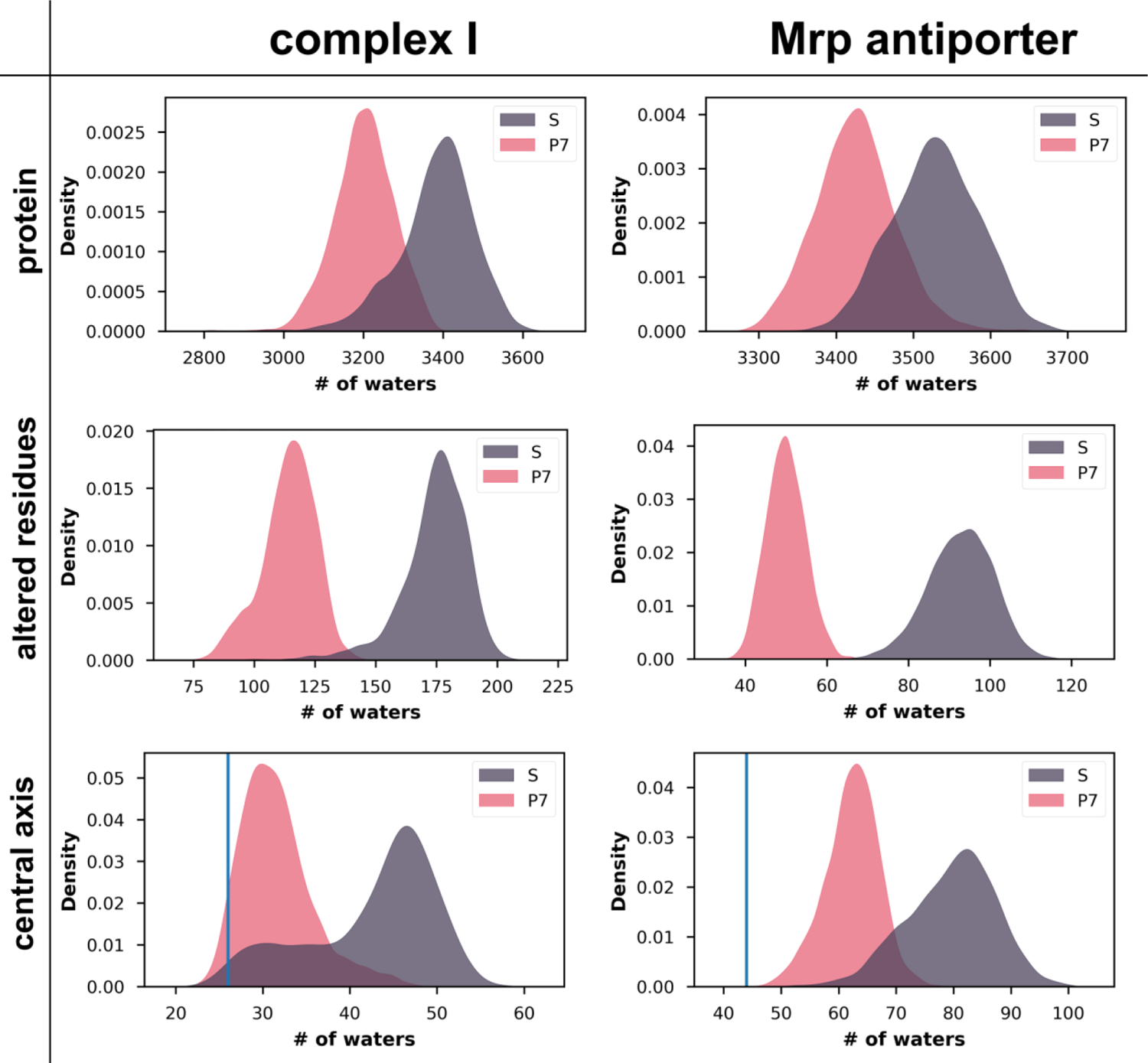
KDE plots for the number of water oxygen atoms in contact within 4 Å of protein residues, with the left column showing data for complex I (core membrane subunits) and the right for Mrp antiporter. Top panels show the number of waters in contact with all protein atoms, middle panels show for protein residues that change protonation state in P7 state simulations, and lower panels show the number of waters in contact with residues in the functionally relevant horizontal axis (Fig. 1). Blue vertical lines in the lower panels indicate the number of waters resolved in the cryo-EM structures that are in contact with the residues in the central hydrophilic axis.

In both proteins, a hydrophilic axis runs through the central part of the membrane domain, which is thought to be the structural basis for proton transfer reactions ^43–48^. We next evaluated the time evolution of internal water molecules in this hydrophilic axis for both states in both proteins. We observed that the number of internal water molecules in the P7 state remains lower and closer to the water content observed in the structures, whereas in S state runs, extensive hydration of the proteins is observed (Fig. 3, lower panel and Fig. S3).

Next, we analyzed the hydrogen bonding between the water molecules and the protein. The time evolution of the hydrogen bonds between protein and structurally resolved water molecules showed a more rapid decline in S states compared to the P7 states, with the latter also displaying a slightly higher number of such interactions in long time scales (Fig. 4). This indicates that although water exchange does take place in the P7 state, the exchange is less rapid and a greater number of hydrogen bonds that existed in the initial structural state are maintained for long time scales. The stable water-protein hydrogen bonds combined with lower hydration in the P7 state also suggests a lower water exchange rate in these simulations. The details of which protein-structural water hydrogen bonds are stabilized in the Mrp antiporter is shown in Table S3, and notably the stabilization of these hydrogen bonds is associated with a decrease in RMSF.

**Fig 4.**
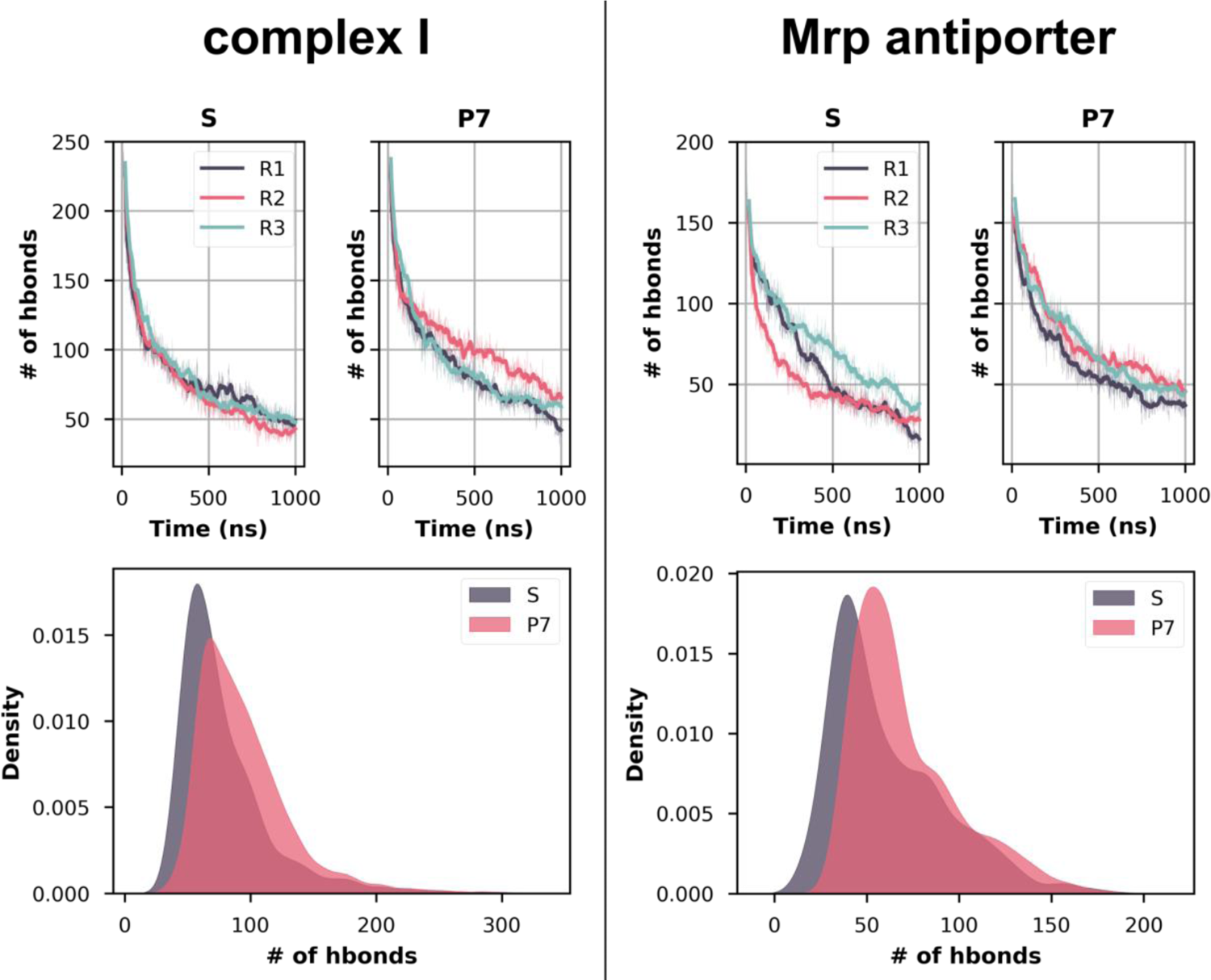
Hydrogen bonds between protein and structural water molecules. (Top) The number of hydrogen bonds throughout the S and P7 simulation trajectories are shown as a time series. The different colors of traces represent different simulation replicas. (Bottom) The data from all trajectories is shown in the lower panels as a density. Only the core membrane subunits of complex I were considered for analysis. The hydrogen bonding distance was cut off at 3.5 Å and the angle at 150.֯

In Mrp antiporter, in the S state most structural waters loose hydrogen bonds with the protein residues in the µs time scale, whereas in P7 state the loss is relatively slower and less extensive in the given time scales (Fig. 4). Interestingly, the loss of hydrogen bonds is much more drastic in complex I simulations, especially in the shorter time scales. To further probe into the kinetics of loss of hydrogen bonding between the structural waters and protein, we fitted the profiles (Fig. 4, upper panels) to exponential function (see methods) and obtained the half-life of protein-water hydrogen bonds (t_1/2_). The results show that on average in complex I P7 state hydrogen bonds survive longer (average t_1/2_ ∼162 ns) compared to S state (average t_1/2_ ∼113 ns). For relatively shorter time scales (ca. 500 ns), a more subtle but similar effect was also observed for Mrp antiporter simulations (average t_1/2_ ∼135 ns in S state compared to ∼145 ns in P7 state).

### Protein-protein interactions

In addition to the protein-water hydrogen bonding interactions, protein-protein hydrogen bonds were also analyzed. The number of protein-protein sidechain hydrogen bonds are consistently found to be higher throughout the P7 state trajectories (Fig. 5). This implies that structural stability is higher in P7 states compared to the S states for both Mrp antiporter and complex I, as also observed in RMSD and RMSF analysis (see above). The lower number of protein-protein interactions is also in agreement with the higher level of hydration observed in the S state. In other words, enhanced water entry/exit in the protein causes protein-protein interactions to be perturbed, resulting in structural destabilization. Specific pairs of residues from the Mrp antiporter that showed a significant increase (>50 %) in hydrogen bonding are shown in Table S4. Interestingly, although only two of the residue pairs undergo changes in protonation state between S and P7 states, still several other pairs show enhancement in hydrogen bonding occupancy, pointing out that long-range effects can occur upon charge change.

**Fig. 5.**
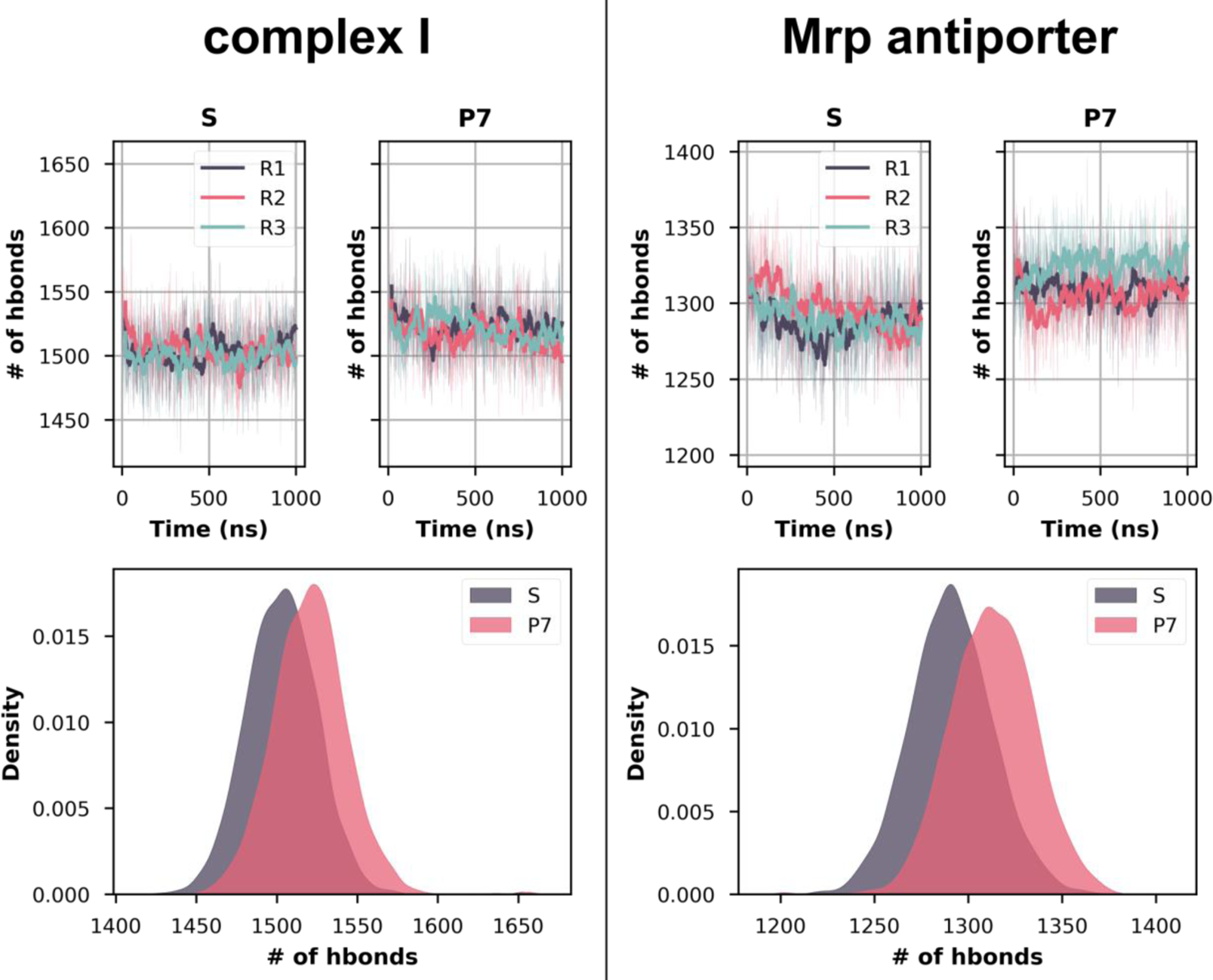
Number of hydrogen bonds between protein sidechains throughout the simulation for complex I and Mrp antiporter. Top panels show the time evolution of all hydrogen bonds for three different simulation replicas, with the hydrogen bonding distance cut off at 3.5 Å and the angle at 150.֯ Bottom panels show the same data as KDE plots. In case of complex I, only core subunits of the membrane arm are considered (see Fig. S4 for full protein).

The above data on protein dynamics and hydration, as well as hydrogen bonding analysis in two different charge states lead to the conclusion that higher protein fluctuations are caused by extensive hydration, which in turn is caused by the highly charged state of the protein, as in the S state. The correct (or near-to-correct) charge state of the protein is thus critical for its structural stability. Hence, prior to launching long time scale MD simulations of high-resolution membrane proteins, a proper charge analysis by means of pKa calculations is recommended.

### Simulations with different pH values

It is clear from the above analysis that the protein simulations that have charge states defined by predetermined protonation states of titratable residues set at pH 7 are well-behaved and are closer to the conformation obtained from cryo-EM. This is true for both global and local mobility, as well as protein hydration. To further explore how altering protonation state affects these characteristics, we performed a series of simulations of the Mrp antiporter complex at 4 more pH values: 5, 6, 8, and 9. The RMSD of the protein (Fig. 6A) was lowest at pH 6 and pH 7, with pH 5 showing the most instability. Similarly, both the number of water molecules in contact with protein (Fig. 6B) and the number of water molecules in the central hydrophilic axis (Fig. 6C) was lowest for pH 6 and pH 7, with pH 5 deviating the most. At pH 5, many of the negatively charged residues become neutral, and positive residues stay positive. Importantly, many histidine residues become doubly protonated, which introduces excess positive charge in several regions of the protein. As a result, bulk water molecules enter the protein, causing instability in protein sidechains and loops, even more than in the S state.

**Fig. 6.**
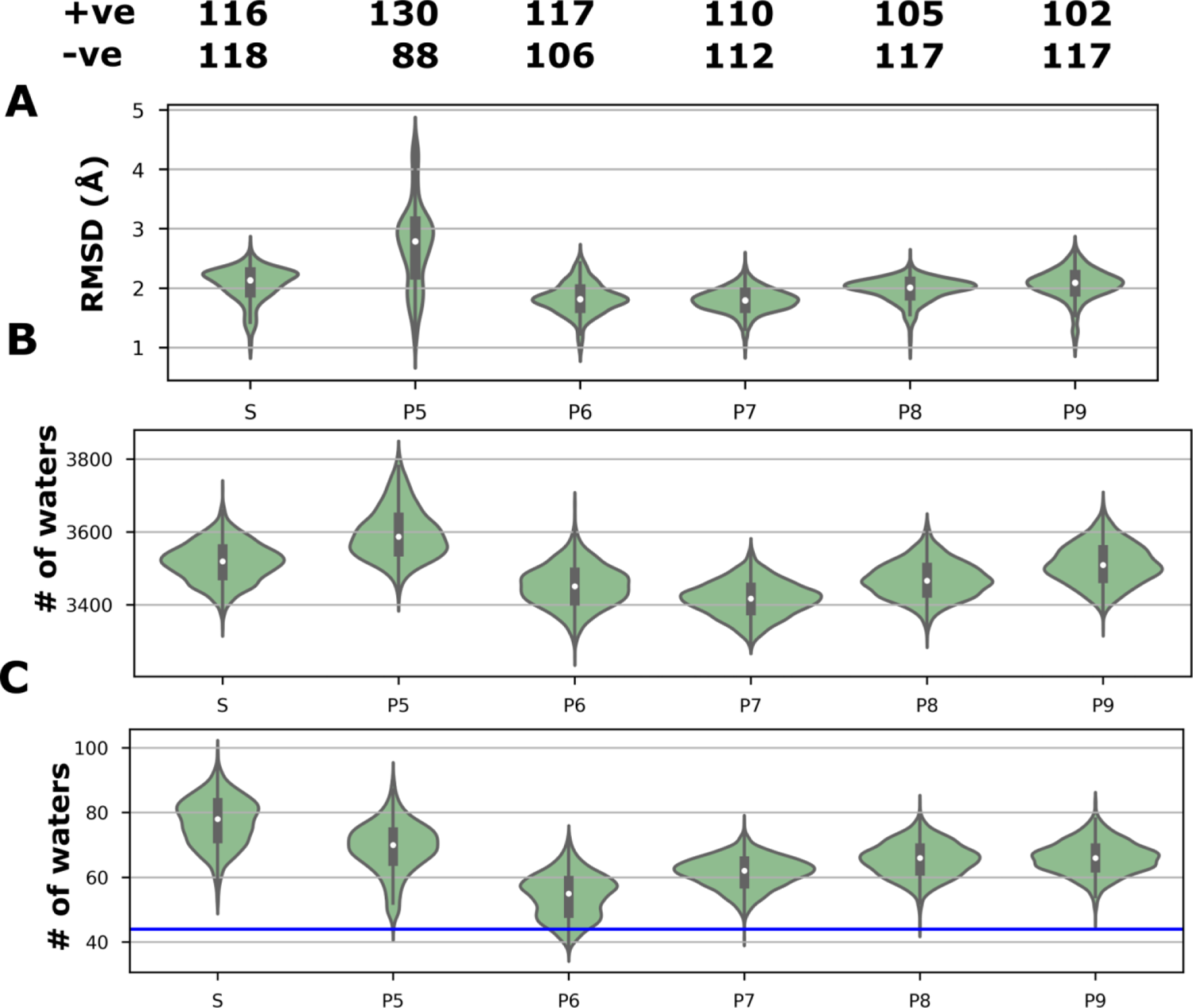
Comparison of Mrp antiporter MD simulations with protonation states set at different pH values. Panel A shows the RMSD of backbone atoms, panel B shows the number of water molecules within 4 Å of all protein residues, and panel C shows the number of waters in the functionally relevant central axis (Fig. 1), with the blue line showing the number of waters in contact with protein observed in the cryo-EM structure. Data is shown as violin plots, where the shaded area is a KDE plot of all trajectory data and in the center is a box plot displaying the median (white dot with the black box indicating the interquartile range). The total number of positively and negatively charged residues at different pH are listed above panel A.

### Charge state assignment based on cryo-EM density

The above analysis highlights the importance of correct charge state description of protein, which is necessary for its stability during MD simulations. However, prediction of charge state by pKa calculations has its limitations. We thus attempted to deduce the protonation states of amino acids from the high resolution cryo-EM data. Cryo-EM density maps can be used to identify the charge states of acidic amino acid residues (glutamates and aspartates), because neutral and negatively charged atoms have vastly different scattering amplitudes ^14, 17^ and thus different cryo-EM density profiles. By carefully analyzing the cryo-EM density in our high-resolution maps, we obtained the charge states of selected glutamate and aspartate residues and compared those to the pKa values obtained based on the modeled structure. We found that there was an agreement between the charge state assignment based on the cryo-EM maps (EMD-12742 for complex I and EMD-14124 for the Mrp antiporter) and the predicted pKa based on the derived atomic models (PDBs 7O71 and 7QRU, respectively). However, several acidic residues showed discrepancies; the pKa calculations predicted some carboxylates to be charge neutral that were assigned to be anionic by the cryo-EM density analysis (Table 1). A possible explanation for this discrepancy is that although the protein model-based pKa predictions are largely correct (especially for buried residues), the positions of the (invisible) sidechains of charged glutamates and aspartates were incorrectly modeled, resulting in overestimated structure-based pKas. To fix this discrepancy between the two estimates, the cryo-EM density map of respiratory complex I (EMD-12742) was revisited. We remodeled the sidechains of selected residues that are predicted to be anionic based on missing cryo-EM densities using Coot ^49^ and re-performed the pKa prediction on the new model. A total of 7 glutamates and aspartates were remodeled one at a time (Table 1).

**Table 1.** Charge state of selected acidic amino acid residues in complex I from cryo-EM density data (EMD 12742) and pKa calculations, based on PDB 7O71 coordinates or on a remodeled carboxylate sidechain. All carboxylates were remodeled independently. ^a^ D67^ND3^ was not remodeled, but instead the new pKa value is based on the structure with E147^ND1^ remodeled.

|  | Density charge estimate | Original pKa | Remodeled pKa |
| --- | --- | --- | --- |
| E147<sup>ND1</sup> | Charged | 9.11 | 7.03 |
| E206<sup>ND1</sup> | Charged | 8.85 | 6.37 |
| E210<sup>ND1</sup> | Charged | 7.96 | 5.69 |
| E231<sup>ND1</sup> | Charged | 7.44 | 6.56 |
| D67<sup>ND3</sup> | Neutral | 7.18 | 9.34<sup>a</sup> |
| E69<sup>ND3</sup> | Charged | 9.27 | 9.16 |
| E30<sup>ND4L</sup> | Charged | 7.59 | 8.69 |
| E66<sup>ND4L</sup> | Charged | 9.81 | 10.16 |

The residues E231^ND1^ and E147^ND1^ of the functionally relevant E-channel in complex I ^43^ were both predicted to be neutral based on pKa calculations but charged based on density analysis. When pKa calculations were performed on re-modeled sidechains, it led to a drop in their sidechain pKa and stabilization of their deprotonated states in good agreement with cryo-EM density-based assignment (Table 1). Interestingly, the remodeled sidechain conformation of E147^ND1^ not only improved the pKa of the residue itself, but also that of residues in its surroundings, e.g. D67^ND3^ (Fig. 7, Table 1). A similar improved agreement between pKa prediction and cryo-EM-based charge state assignment was also observed for E206 and E210 from the ND1 subunit (Table 1), both of which are known to be central for protein function^43^. However, pKa predictions of E30 and E66 from ND4L, as well as E69 from ND3 could not be improved despite their remodeled sidechains. One potential reason for this could be that the surroundings in two different conformations do not differ as drastically as in the other cases. Furthermore, the approximate nature of pKa predictions is well known, and with no explicit treatment of neighboring solvation can result in the overestimation of pKa values. All in all, remodeling of several sidechains not only improved the charged state assignment, but also improved the atomic modeling of cryo-EM density data (see discussion).

**Fig 7.**
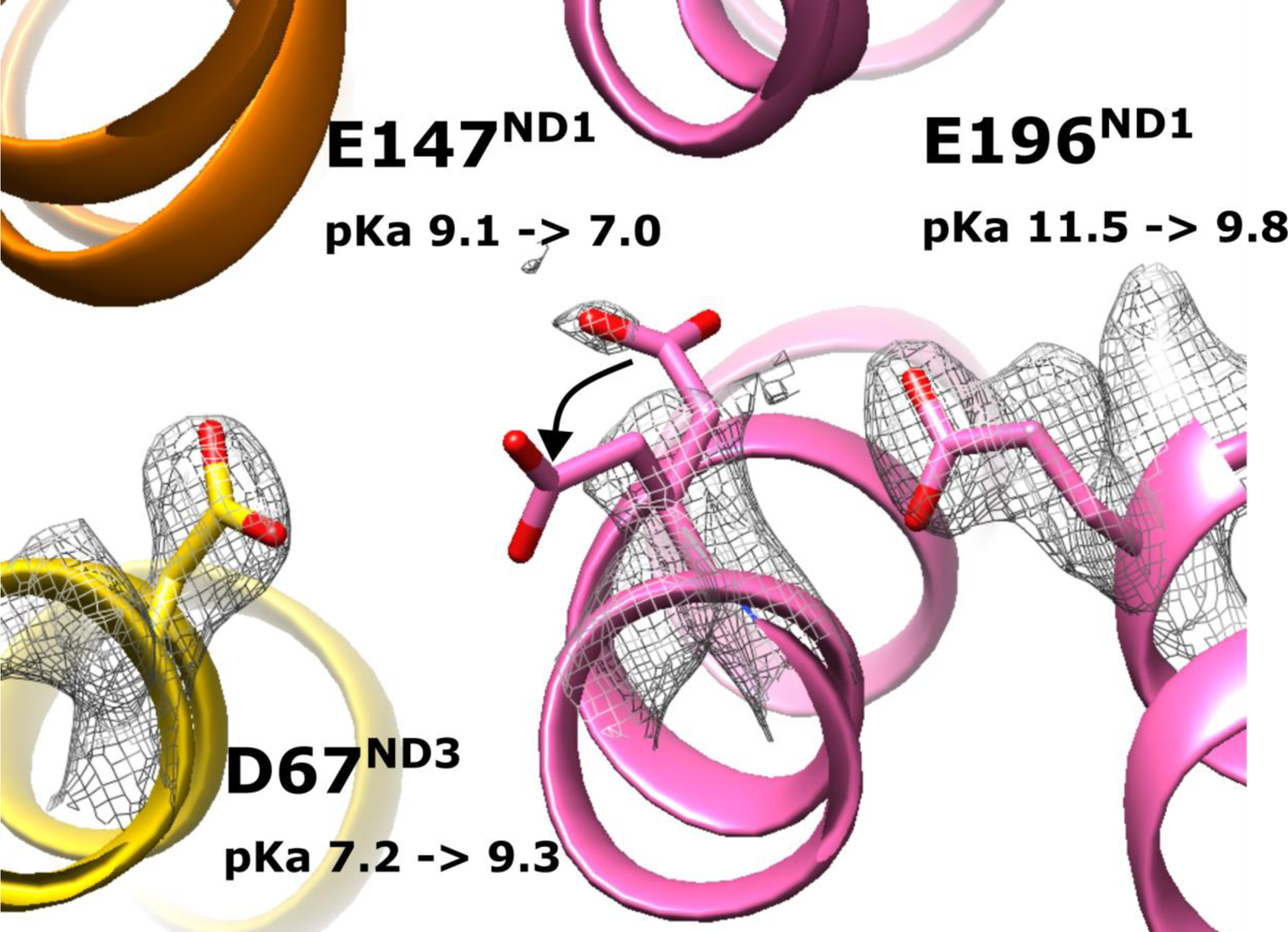
The remodeling of the sidechain of E147^ND1^ (indicated by black arrow) improved its pKa as well as that of neighboring D67^ND3^. The first value of pKa is based on the original coordinates (PDB 7O71), while the second value is after remodeling the sidechain of E147^ND1^. The cryo-EM density of the three residues is shown as grey mesh.

## Discussion

Protonation states of amino acids play a key role in protein structure stability and function. However, it is extremely challenging to obtain an accurate estimate of the pKa of an amino acid both by experiments and computations, and as a result in most cases, pKa values and how they change during enzymatic action remains unknown. Because of this, MD simulations are often performed by modeling amino acids in their fixed protonation states, including standard charged states. Here we show that MD simulations of a membrane protein with all amino acids modeled in their standard charged states results in lower stability of the protein overall and drives the conformation of the protein far from its starting point that is the cryo-EM conformation. Instead, if MD simulation is performed by fixing the charged states of titratable amino acids based on pKa calculations, the conformational state remains closer to the original structure. Our analysis of protein mobility and hydration, as well as residue-water and residue-residue interactions supports this notion. Also, since the structural conformation is retained in MD simulations performed after pKa calculations, we propose these more closely match the charge states of amino acids that existed during the cryo-EM experiment.

Our results also indicate that the relationship between protein stability and hydration is strong, and that reduced hydration leads to a more stable simulation system which can maintain important interactions. When leaving all protonatable residues in their standard charged states, hydration is considerably enhanced, resulting in important protein-protein interactions being broken, thereby reducing stability.

The higher level of hydration observed in standard state simulations is primarily due to the charged states of buried titratable amino acid residues. These charged amino acid residues in the low protein dielectric interior create an unfavorable high-energy scenario due to poor solvation. This establishes an electric field, pulling bulk water molecules into the protein and enhancing solvation similar to the nanoscopic electroporation described earlier ^50^, see also ^33^. We also note that protein hydration obtained from MD simulations performed in pre-defined charge state is much closer to the structural hydration than that from the MD simulations performed in standard state of the residues. Despite this, the hydration obtained from MD simulations is in general higher than the hydration observed in the structure. The reason for this discrepancy is in part due to the underestimation of protein hydration in the cryo-EM maps, where only highly occupied and tightly bound water molecules can be observed.

By performing MD simulations in pre-determined charge state from pKa calculations, one not only investigates the state that was captured during structure resolution, but also its time evolution more accurately. This is particularly useful in situations where there are a particularly large number of titratable residues present in the protein with possible functional relevance, such as in photosynthetic and respiratory enzymes, as well as proteins that couple translocation of ions or metabolites to proton transfer reactions. However, empirical pKa calculations or more thorough continuum electrostatics based pKa estimations have limited accuracy and site-site interaction energy terms can often dominate, leading to large scale pKa shifts with subtle changes in structure. Thus, it is important to use an accurate input model, based on high-resolution data where there is less uncertainty in sidechain placements. Below ∼2.5 Å resolution the correct rotamers of most sidechains can be confidently modeled, with the key exception being negatively charged sidechains in cryo-EM maps, which have negative atomic scattering factors ^13, 17, 18^ and thus are usually invisible beyond the Cβ atoms. As modeling and refinement programs do not take this into account, the sidechain orientation of these residues is unreliable ^18^. On the other hand, this fact makes it possible to deduce the charge state of acidic residues from the cryo-EM density.

Here we propose a method that may assist to improve modeling of sidechains of acidic residues. By performing fast empirical pKa calculations on the 3D model, pKas of all acidic residues can be estimated. If the predicted pKa value suggests a neutral state of an acidic residue that is expected to be anionic based on density analysis, this warrants remodeling of the sidechain of the amino acid. If the remodeled sidechain conformation yields a lower pKa, this is likely to be the more accurate conformation. By combining pKa calculations with sidechain remodeling on selected acidic residues of complex I, we show here that this is feasible. In our future work, we aim to automate this process and obtain more accurate models of sidechains of acidic amino acid residues from the cryo-EM data.

## Methods

We performed all-atom molecular dynamics simulations of complex I from *Yarrowia lipolytica* (PDB: 7O71)^43^ and Mrp antiporter from *Bacillus pseudofirmus* (PDB: 7QRU, altloc B)^44^. Both proteins were placed in a mixed lipid bilayer and solvated with TIP3P and NaCl. Gromacs versions 2020 and 2021 ^51^ were used for minimization, equilibration, and production runs. Full details of the molecular dynamics set ups can be found in previous work on complex I^43^ and Mrp antiporter^44^. The production runs were performed without any constraints in an NPT ensemble, using Nosé–Hoover thermostat^52, 53^ at 310 K and Parrinello-Rahman barostat^54^ at 1 atm. We employed the LINCS algorithm to achieve a 2 fs timestep^55^, and the particle-mesh Ewald method^56^ with a cut-off of 12 Å to handle electrostatic interactions. The cut-off for van der Waals interactions was 12 Å, with a switching distance of 10 Å. Table S5 lists the simulations setups, and the S and P7 states for complex I and Mrp antiporter are extended simulations from previous work. The P5, P6, P8, and P9 simulations of Mrp are new systems constructed using the same protocol.

Estimates of pKa were performed using the Propka software package^39^. The calculations were performed on the PDB structures of both complex I and Mrp. Asp, Glu and His were considered protonated (neutral) if their pKa was more than 7 in the P7 simulations, while Lys was considered deprotonated if its pKa was less than 7. The same process was performed in the P5, P6, P8, and P9 setups, with their respective pKa cutoffs changed. His was considered with the δ-nitrogen protonated when neutral.

Exponential fit of hydrogen bond decay was calculated using the SciPy *curve_fit* function^57^. We used a mono-exponential of the form *y = me^(−tx)^ + b*, where m, t, and b were parameters that could vary. The quality of the fit was determined using R², obtaining values > 0.9. The half-life was calculated as *t*_1/2_ = t × ln 2.

The electron density analysis and remodeling of the resolved structures was done using Coot^49^. The change in the glutamic and aspartic acid conformations was achieved with the rotamers function in Coot. The key requirement for the new position of the sidechain was a substantial deviation from its previous position, as well as avoiding clashes with its surroundings. Care was taken not to position the side chain in positive density. Visualization, analysis, and figure preparation was performed using VMD, Pymol and UCSF Chimera ^58^.

## Acknowledgements

VS acknowledges research funding from the Academy of Finland, the Jane and Aatos Erkko Foundation, the Sigrid Juselius Foundation and the Magnus Ehrnrooth Foundation. JV and VZ were supported by the German Research Foundation (CRC 1507 – Membrane-associated Protein Assemblies, Machineries, and Supercomplexes; P14). High-performance computing time from Center for Scientific Computing, Finland is acknowledged. AD acknowledges travel grant support from the Finnish Concordia Fund and the Magnus Ehrnrooth foundation. We also thank Dr. Outi Haapanen for simulation data of Mrp antiporter.

