## Supplementary File for "Role of protonation states in stability of molecular dynamics simulations of high-resolution membrane protein structures"

### Supplementary Information

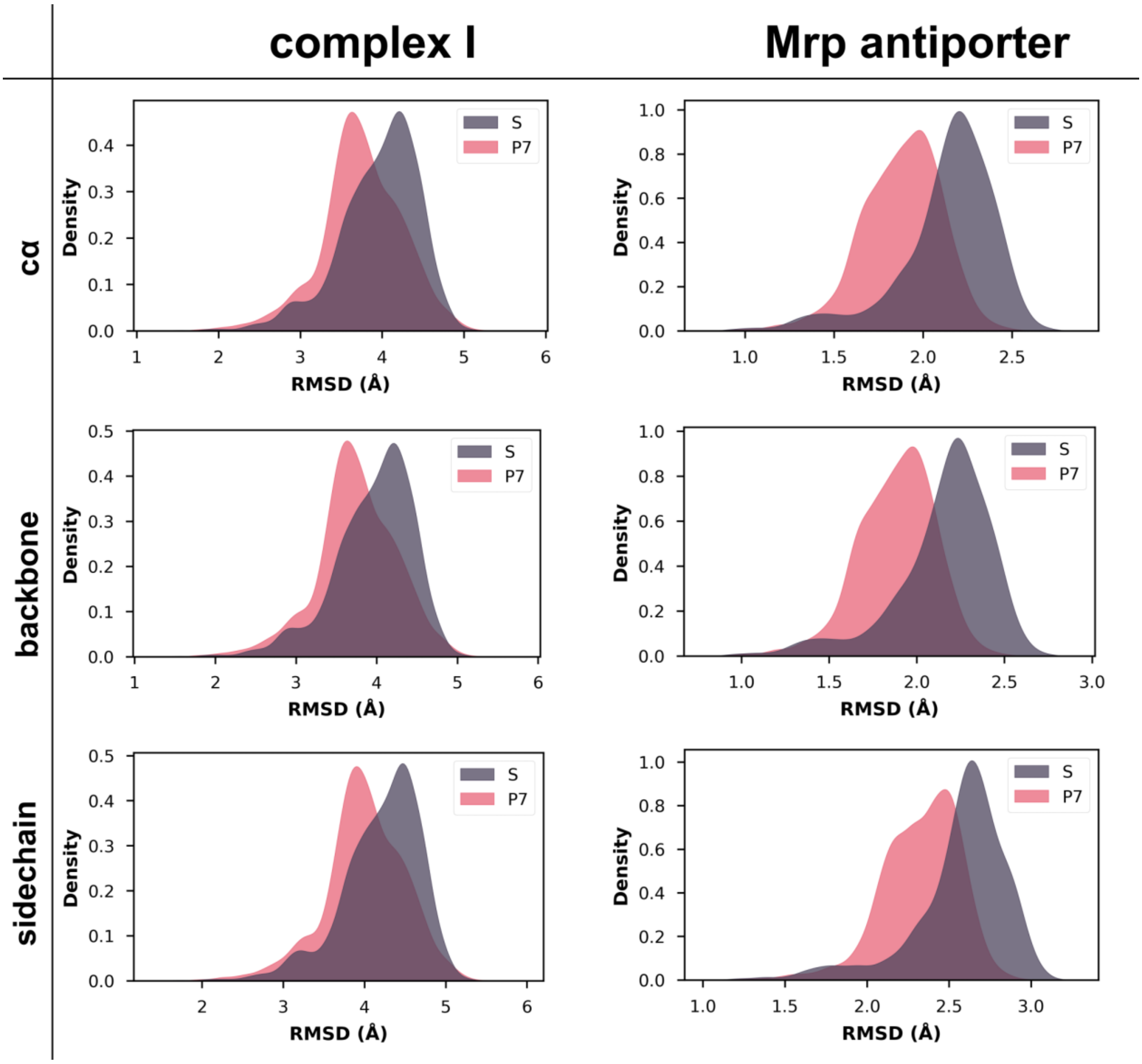

**Fig S1 –** RMSD (root mean square deviation) over time for both complex I and Mrp systems. The data is combined for all simulations replicas (3x1000ns, see methods), and the RMSD is measured every 1 ns. The charts show the distribution of values in the S and P7 states using a kernel density estimate function (KDE) with combined data of all three replicas. Upper panels show the RMSD for Cα atoms, middle panels for backbone atoms, and lower panels for all protein sidechain atoms excluding hydrogens. Atoms in the hydrophilic domain of complex I were included in all plots.

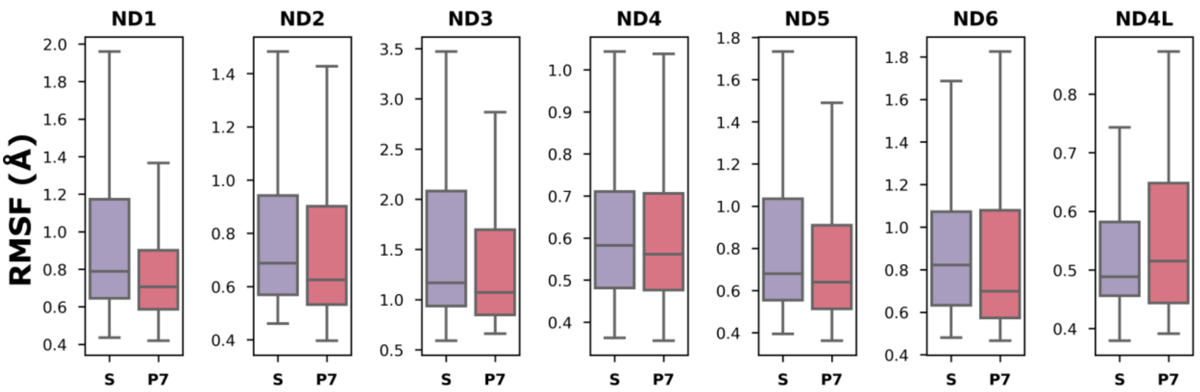

**Fig. S2** - RMSF (root mean square fluctuation) of Cα atoms of core membrane-bound complex I subunits. The data shown is based all simulation data for S and P7 states (3 x 1 µs, see methods). The shaded box represents the interquartile range, with the middle line showing the median. The upper and lower lines are the maximum and minimum values, respectively.

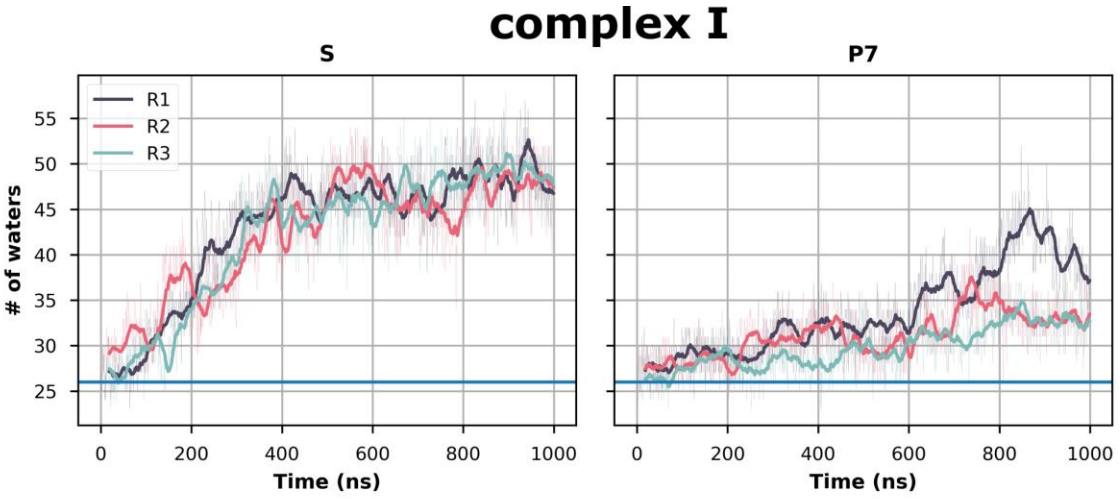

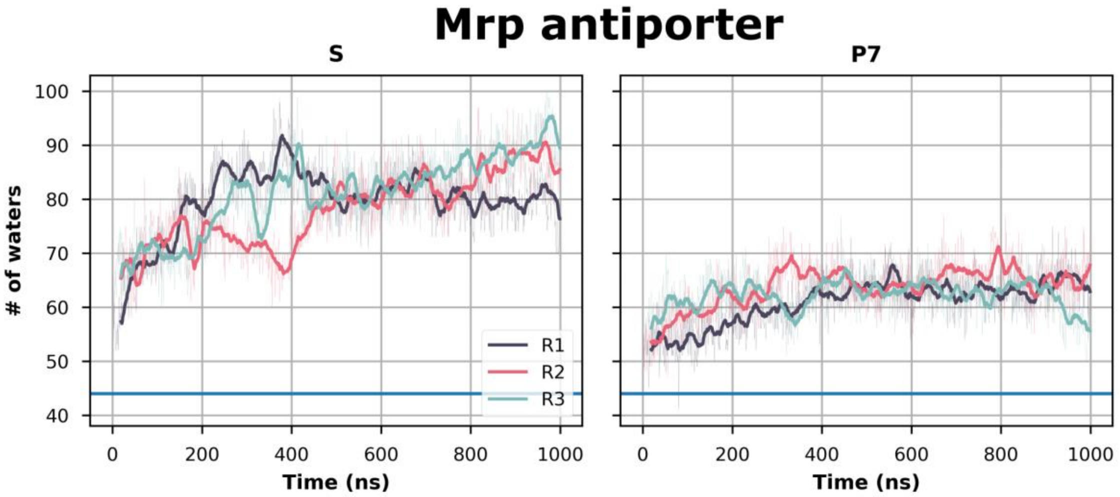

**Fig. S3** – Time-series of hydration in the hydrophilic central axis of complex I (upper) and Mrp antiporter (lower). Water residues were counted every frame within 4 Å of selected residues. The blue horizontal line represents the number of water molecules seen in the cryo EM structure.

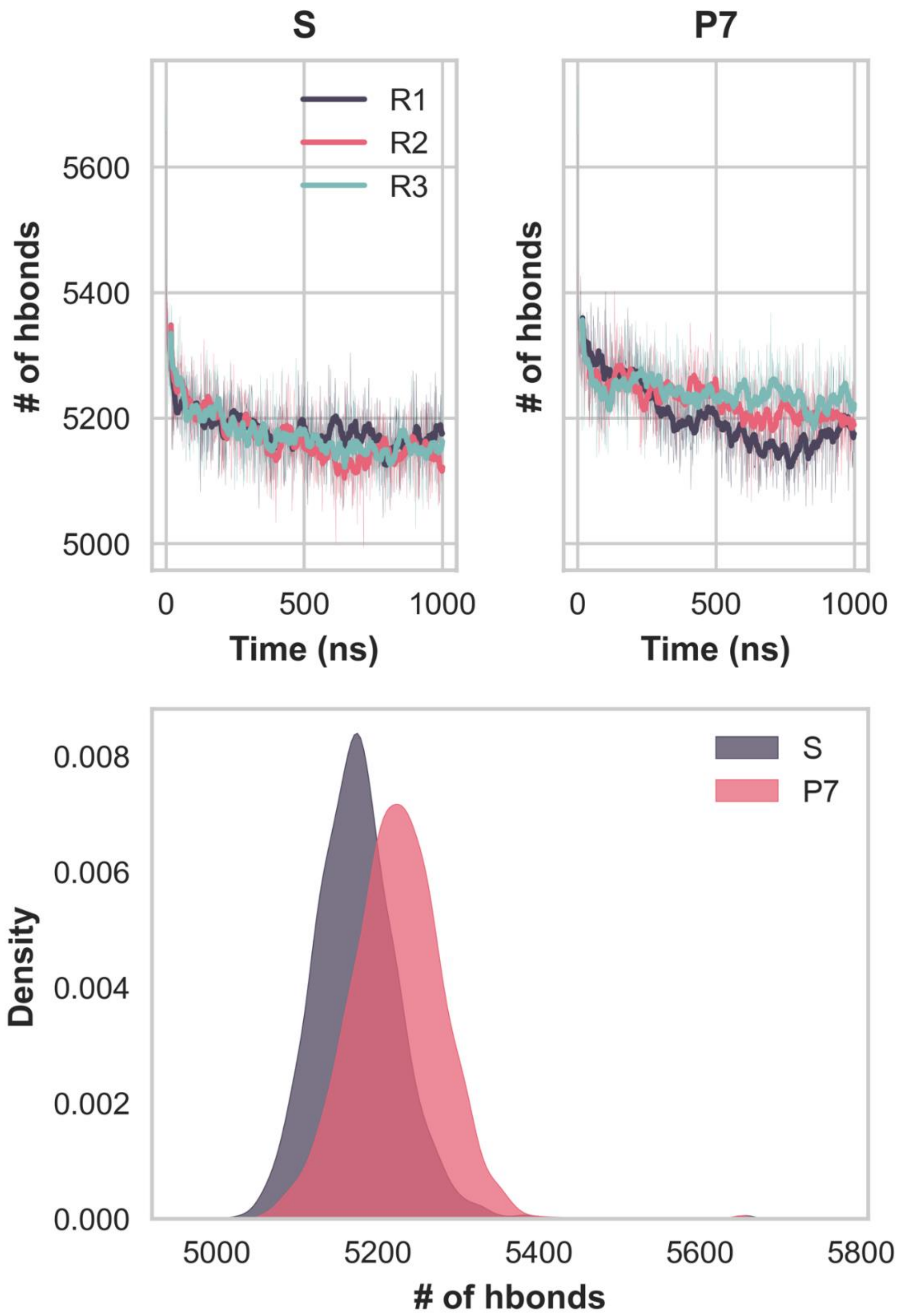

**Fig. S4** - Number of hydrogen bonds between protein sidechains throughout the simulation for all protein subunits in complex I. Top panels show the time evolution of all hydrogen bonds for three different simulation replicas, with the hydrogen bonding distance was cut off at 3.5 Å and the angle at 150̊. Bottom panels show the same data as KDE plots.

**Table S1 –** Titratable residues in complex I which change protonation state in setup P7. The pKa calculated by PropKa is shown. Only residues in the core membrane subunits are shown.

| **Subunit** | **Residue** | **pKa** | **P7** |
|---|---|---|---|
| ND1 | Glu147 | 11.00 | 0 |
| | Glu196 | 9.52 | 0 |
| | Glu206 | 8.96 | 0 |
| | Glu210 | 7.41 | 0 |
| | Glu231 | 7.49 | 0 |
| | Lys285 | 6.61 | 0 |
| ND2 | Asp39 | 7.97 | 0 |
| | Asp69 | 8.43 | 0 |
| | Lys282 | 6.27 | 0 |
| ND3 | Asp67 | 7.19 | 0 |
| | Glu39 | 7.04 | 0 |
| | Glu69 | 9-27 | 0 |
| ND4 | Glu142 | 8.16 | 0 |
| | Glu395 | 7.57 | 0 |
| | Lys252 | 6.75 | 0 |
| | Lys299 | 6.47 | 0 |
| ND5 | Asp178 | 9.04 | 0 |
| | Asp397 | 7.91 | 0 |
| | Glu80 | 7.35 | 0 |
| | Glu503 | 8.79 | 0 |
| | Lys339 | 6.48 | 0 |
| | Lys581 | 6..80 | 0 |
| ND4L | Asp49 | 7.20 | 0 |
| | Glu30 | 7.60 | 0 |
| | Glu66 | 9.77 | 0 |

**Table S2 –** Titratable residues in Mrp antiporter whose charge states were changed in MD setups. The setups P5-9 represent pH values of 5-9. The pKa estimate based on PropKa calculation is shown.

| Subunit | Residue | pKa | P5 | P6 | P7 | P8 | P9 |
|---|---|---|---|---|---|---|---|
| MrpA | Asp678 | 7.36 | 0 | 0 | 0 | -1 | -1 |
| | Asp771 | 7.65 | 0 | 0 | 0 | -1 | -1 |
| | Asp776 | 5.77 | 0 | -1 | -1 | -1 | -1 |
| | Glu59 | 6.28 | 0 | 0 | -1 | -1- | -1 |
| | Glu107 | 5.51 | 0 | -1 | -1 | -1 | -1 |
| | Glu140 | 6.95 | 0 | 0 | -1 | -1 | -1 |
| | Glu157 | 5.37 | 0 | -1 | -1 | -1 | -1 |
| | Glu366 | 6.50 | 0 | 0 | -1 | -1 | -1 |
| | Glu409 | 6.79 | 0 | 0 | -1 | -1 | -1 |
| | Glu503 | 5.59 | 0 | -1 | -1 | -1 | -1 |
| | Glu687 | 7.75 | 0 | 0 | 0 | -1 | -1 |
| | Glu707 | 5.90 | 0 | -1 | -1 | -1 | -1 |
| | Glu780 | 9.52 | 0 | 0 | 0 | 0 | 0 |
| | His5 | 5.13 | +1 | 0 | 0 | 0 | 0 |
| | His60 | 5.89 | +1 | 0 | 0 | 0 | 0 |
| | His155 | 5.77 | +1 | 0 | 0 | 0 | 0 |
| | His230 | 5.84 | +1 | 0 | 0 | 0 | 0 |
| | His365 | 5.57 | +1 | 0 | 0 | 0 | 0 |
| | His470 | 7.17 | +1 | +1 | +1 | 0 | 0 |
| | His528 | 5.89 | +1 | 0 | 0 | 0 | 0 |
| | Lys223 | 6.65 | +1 | +1 | 0 | 0 | 0 |
| | Lys254 | 5.90 | +1 | 0 | 0 | 0 | 0 |
| | Lys299 | 6.84 | +1 | +1 | 0 | 0 | 0 |
| | Lys353 | 6.43 | +1 | +1 | 0 | 0 | 0 |
| | Lys408 | 6.37 | +1 | +1 | 0 | 0 | 0 |
| MrpB | Asp121 | 7.35 | 0 | 0 | 0 | -1 | -1 |
| | Asp142 | 5.53 | 0 | -1 | -1 | -1 | -1 |
| | Glu113 | 6.49 | 0 | 0 | -1 | -1 | -1 |
| | Glu141 | 5.83 | 0 | -1 | -1 | -1 | -1 |
| MrpC | Asp111 | 5.39 | 0 | -1 | -1 | -1 | -1 |
| | His58 | 5.3 | +1 | 0 | 0 | 0 | 0 |
| | His98 | 5.06 | +1 | 0 | 0 | 0 | 0 |
| MrpD | Glu60 | 5.69 | 0 | -1 | -1 | -1 | -1 |
| | Glu137 | 6.55 | 0 | 0 | -1 | -1 | -1 |
| | Glu467 | 5.10 | 0 | -1 | -1 | -1 | -1 |

| | | | | | | | |
|---|---|---|---|---|---|---|---|
| | His26 | 5.67 | +1 | 0 | 0 | 0 | 0 |
| | His266 | 5.50 | +1 | 0 | 0 | 0 | 0 |
| | His366 | 5.08 | +1 | 0 | 0 | 0 | 0 |
| | Lys161 | 8.87 | +1 | +1 | +1 | +1 | 0 |
| | Lys219 | 8.01 | +1 | +1 | +1 | +1 | 0 |
| | Lys250 | 6.31 | +1 | +1 | 0 | 0 | 0 |
| | Lys297 | 7.88 | +1 | +1 | +1 | 0 | 0 |
| | Lys337 | 5.61 | +1 | 0 | 0 | 0 | 0 |
| | Lys392 | 7.38 | +1 | +1 | +1 | 0 | 0 |
| | Lys424 | 7.05 | +1 | +1 | +1 | 0 | 0 |
| MrpE | Asp74 | 5.32 | 0 | -1 | -1 | -1 | -1 |
| | Glu67 | 5.72 | 0 | -1 | -1 | -1 | -1 |
| | Glu150 | 5.77 | 0 | -1 | -1 | -1 | -1 |
| | His146 | 6.88 | +1 | +1 | 0 | 0 | 0 |
| MrpF | Asp38 | 7.04 | 0 | 0 | 0 | -1 | -1 |
| | Glu56 | 5.50 | 0 | -1 | -1 | -1 | -1 |
| | Glu62 | 5.10 | 0 | -1 | -1 | -1 | -1 |
| | Lys80 | 8.84 | +1 | +1 | +1 | +1 | 0 |
| MrpG | Asp31 | 5.56 | 0 | -1 | -1 | -1 | -1 |
| | Glu64 | 5.49 | 0 | -1 | -1 | -1 | -1 |
| | His62 | 6.25 | +1 | +1 | 0 | 0 | 0 |
| | Lys41 | 7.49 | +1 | +1 | +1 | 0 | 0 |

**Table S3 –** Occupancy of hydrogen bond between structural water and protein sidechain in MD simulations of Mrp antiporter. The subunit names shown in superscript (A/D/F – MrpA/MrpD/MrpF). The RMSF of the residue sidechain is also shown. The data is calculated based on all simulation frames from all trajectories. RMSF is shown as an average of all heavy sidechain atoms. Hydrogen bonds which stabilized more than 20% are shown here. Note complex I had > 90 such hydrogen bonds that stabilized more than 20% occupancy.

| | **Hydrogen bond occupancy (%)** | | | **Sidechain RMSF (Å)** | | |
|---|---|---|---|---|---|---|
| | S | P | Change | S | P | Change |
| **T264$^{A}$** | 11 | 61 | +50 | 0.83 | 0.74 | -0.09 |
| **N763$^{A}$** | 6 | 48 | +42 | 0.68 | 0.66 | -0.02 |
| **S479$^{A}$** | 5 | 41 | +36 | 1.01 | 0.89 | -0.12 |
| **D128$^{D}$** | 8 | 34 | +26 | 0.92 | 0.79 | -0.13 |
| **Y258$^{A}$** | 0 | 24 | +24 | 0.70 | 0.63 | -0.04 |
| **E62$^{F}$** | 10 | 31 | +21 | 1.03 | 0.96 | -0.08 |

**Table S4 –** Hydrogen bond occupancy between selected protein sidechains in Mrp antiporter. The data is calculated based on all simulation frames from all trajectories. The asterisk (*) denotes residues that were neutralized in the P state. Interactions present in the cryo-EM structure are in green.

| | **Hydrogen bond occupancy (%)** | | |
|---|---|---|---|
| | S | P | Change |
| **H349$^{A}$-K254$^{A}$*** | 0 | 76 | +76 |
| **T306$^{A}$-H349$^{A}$** | 0 | 66 | +66 |
| **S421$^{D}$-H303$^{D}$** | 18 | 82 | +64 |
| **Y447$^{A}$-S305$^{A}$** | 7 | 71 | +64 |
| **Y161$^{A}$-D572$^{A}$** | 1 | 47 | +56 |
| **K165$^{A}$-D572$^{A}$** | 12 | 67 | +55 |
| **S170$^{D}$-E137$^{D}$** | 32 | 86 | +54 |
| **H333$^{D}$-K250$^{D}$*** | 0 | 52 | +52 |
| **R194$^{A}$-E59$^{A}$** | 32 | 82 | +51 |

**Table S5 –** List of simulation setups and lengths.

| | State | Replicas x length |
|---|---|---|
| **Complex I** | S | 3 x 1000ns |
| | P7 | 3 x 1000ns |
| **Mrp antiporter** | S | 3 x 1000ns |
| | P7 | 3 x 1000ns |
| | P5 | 3 x 645ns |
| | P6 | 3 x 645ns |
| | P8 | 3 x 645ns |
| | P9 | 3 x 645ns |
